# A Cell type-specific Class of Chromatin Loops Anchored at Large DNA Methylation Nadirs

**DOI:** 10.1101/212928

**Authors:** Mira Jeong, Xingfan Huang, Xiaotian Zhang, Jianzhong Su, Muhammad S. Shamim, Ivan D. Bochkov, Jaime Reyes, Haiyoung Jung, Emily Heikamp, Aviva Presser Aiden, Wei Li, Erez Lieberman Aiden, Margaret A. Goodell

## Abstract

Higher order chromatin structure and DNA methylation are implicated in multiple developmental processes, but their relationship to cell state is unknown. Here, we found that large (~10kb) DNA methylation nadirs can form long loops connecting anchor loci that may be dozens of megabases apart, as well as interchromosomal links. The interacting loci comprise ~3.5Mb of the human genome. The data are more consistent with the formation of these loops by phase separation of the interacting loci to form a genomic subcompartment, rather than with CTCF-mediated extrusion. Interestingly, unlike previously characterized genomic subcompartments, this subcompartment is only present in particular cell types, such as stem and progenitor cells. Further, we identify one particular loop anchor that is functionally associated with maintenance of the hematopoietic stem cell state. Our work reveals that H3K27me3-marked large DNA methylation nadirs represent a novel set of very long-range loops and links associated with cellular identity.

**Summary:** Hi-C and DNA methylation analyses reveal novel chromatin loops between distant sites implicated in stem and progenitor cell function.

## TEXT

In the human genome, cytosine residues located in CpG dinucleotides are often, but not always, methylated (5-methyl-C). CpG islands – genomic intervals, typically 300-3000bp in length, containing many CpG dinucleotides – are an important exception (*1*). Frequently located near promoters, CpG islands are typically unmethylated when the nearby gene is active. Yet, despite extensive study, the mechanisms that underlie the relationship between DNA methylation and gene transcription are poorly understood.

One possibility is that the absence of DNA methylation leads to changes in 3d chromatin architecture that influence transcription. In recent years, experiments combining DNA-DNA proximity ligation with high-throughput sequencing (Hi-C) have made it possible to generate high-resolution maps of chromatin architecture by measuring the frequency of contact between all pairs of loci, genome-wide (*2–6*). These experiments – whose results are typically represented as a heatmap in which every pixel indicates the contact frequency between a pair or loci – have revealed two mechanisms of chromatin folding. The first is associated with the formation of a class of loops between sites bound by cohesin and CTCF, such that the CTCF motifs lie in the convergent orientation (i.e., they point toward one another) (*7, 8*). To explain this phenomenon, it has been hypothesized that cohesin initially forms small loops between nearby sites, which grow larger through a process of extrusion until an inward-pointing CTCF is encountered (*7–11*). The second mechanism is compartmentalization: the tendency of genomic intervals with similar histone modifications to co-segregate in 3D inside the nucleus (*3, 4*).

We were interested in exploring a potential relationship between DNA methylation and genome architecture, but the typical CpG island is too short to be reliably interrogated by Hi-C, preventing the exploration of these features. However, we recently identified exceptionally long genomic intervals (~3.5-25 Kb) that exhibit low levels of cytosine methylation, dubbed “DNA methylation canyons” (*12, 13*). Canyons, which often contain multiple CpG islands, are strongly preserved across cell types and species. In any given cell, particular canyons are typically either repressed, and decorated with H3K27me3, or active, and decorated with H3K4me3 and H3K27 acetylation (*12*).

Because of their unusual size, methylation canyons are a natural system for exploring the influence of DNA methylation on genome architecture. Because the DNA methyltransferases regulating canyon size are highly expressed in hematopoietic stem/progenitor cells (HSPC) and important for their proper function (*12, 14*), we began by exploring the 3D architecture of HSPCs, performing *in situ* Hi-C experiments at 10kb resolution. Strikingly, we observe the formation of hundreds of long-range loops between large, repressed canyons lying on the same chromosome, as well as evidence for links between canyons lying on different chromosomes. Taken together, our data are consistent with the formation of a subcompartment in which large, repressed DNA methylation canyons from across the genome tend to co-segregate. We show that these features are present, albeit much weaker, after HSPC differentiation and in other differentiated cell types. Our findings indicate that DNA methylation works in tandem with histone modifications to influence the 3D architecture of the human genome.

We began by isolating HSPCs from human umbilical cord blood (UCB) cells using fluorescence-activated cell sorting (FACS), gating for live, lineage negative, CD34+ CD38-cells (Fig 1A, FigS1a). We then generated an *in situ* Hi-C library (*3*), sequenced ~1 billion Hi-C reads (613M contacts) (Table S1), and processed the data using Juicer (*15*), as previously described.

**Figure 1.**
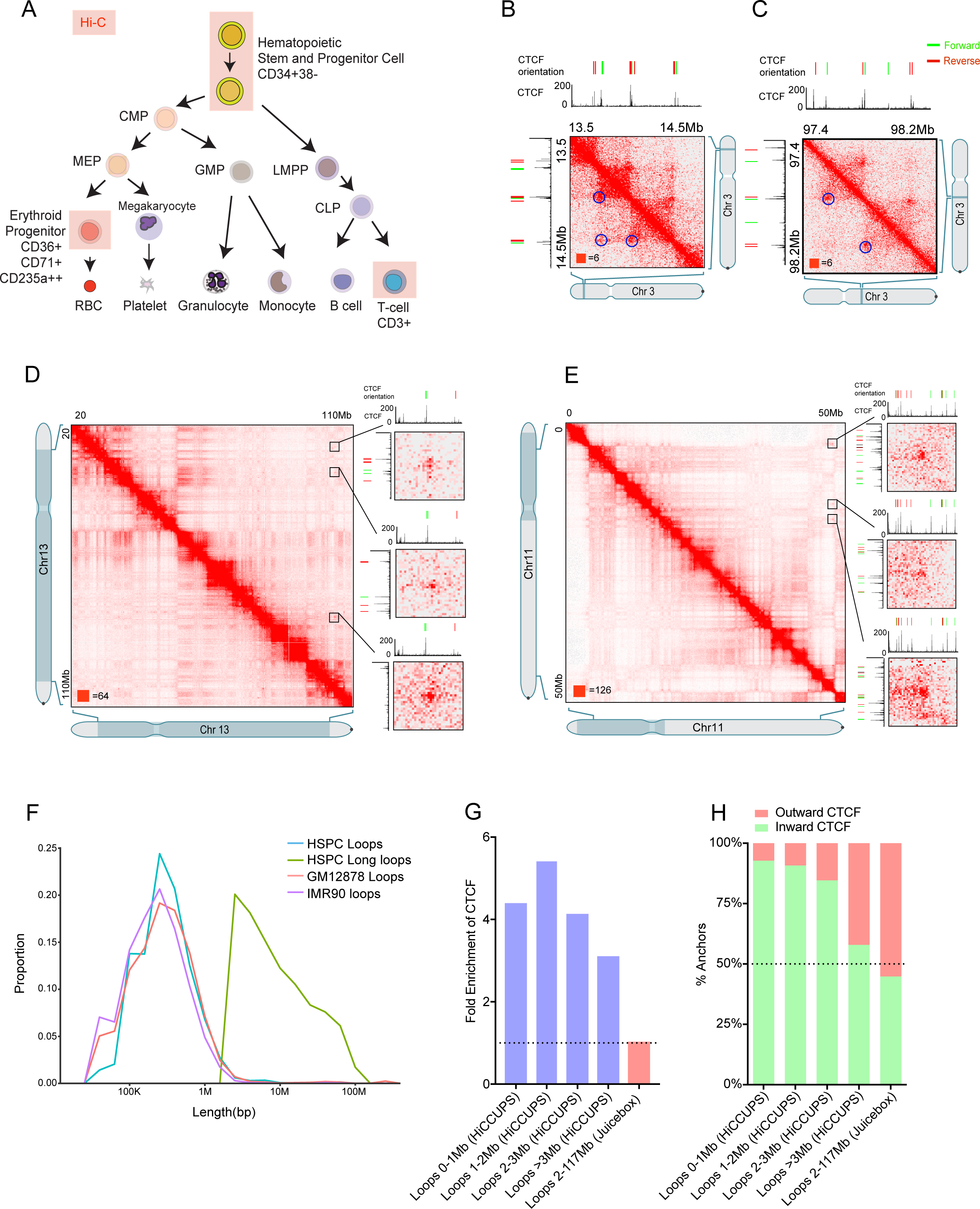
Very long-range interactions in the 3D HSPC genome. **(A)** Diagram of the hematopoietic hierarchy. HSPC, T-cell and Erythroid Progenitors (EPs) were selected for Hi-C profiling (Red shaded box-Hi-C profiling population) **(B)** Example of regular HiCCUPS loop with convergent CTCF motifs on chromosome 3 in the *WNT7A* region at 5kb resolution. **(C)** Example of regular HiCCUPS loop with convergent CTCF motifs on chromosome 3 in the GABRR3 region at 5kb resolution. **(D)** Example of intra multiple long-range interaction on chromosome 13. The matrices are shown at 100kb resolution and blowout of *CDX2*, *NBEA*, *POU4F1* and *ZIC2* region at 10kb and 5kb resolution. **(E)** Example of intra multiple long-range interaction on chromosome 11. The matrices are shown at 100kb resolution and blowout of *FNBP4*, *PSMA1* and *NCR3LG1* region at 10kb and 5kb resolution. **(F)** Length distribution of HiCCUPS loops (blue line) versus Long loops (green line), and loops identified in GM12878 (orange) and IMR90 (purple) cells. **(G)** Fold Enrichment of CTCF binding sites on loop anchors as compared to random translational control regions. (HiCCUPS loops - blue bars, Long loops – red bar) **(H)** Inward and outward orientation of CTCF motifs on loop anchors.

Our loop-calling algorithm (HiCCUPS (*3, 15*) identified 2683 loops in HSPCs, each connecting a pair of loop anchor points on the same chromosome. Many of these loops overlapped with loops that had previously been reported by using *in situ* Hi-C in other cell types (Fig 1B,C, Fig S1b-e). For instance, 2014 of these 2683 loops overlapped the 9448 loops we reported in GM12878 lymphoblastoid cells, and 1832 overlapped the 8040 loops we reported in IMR90 lung fibroblasts. The loop anchors also exhibited similar CTCF-binding profiles to loops reported in previous studies. They were usually bound by CTCF (as assayed by ChIP-Seq: 74%, 4.1-fold enriched vs. random control loci of similar length), with the motifs in the convergent orientation (for 2534 of the 2744 loop anchors with a unique CTCF motif, the motif points inward, 92%, p=6.07×10^−506^). These observations confirmed the accuracy of our Hi-C maps and feature calls in HSPCs.

Interestingly, the CTCF-binding profile of the loop anchors depended on the size of the loop (i.e., how far apart the two loop anchor loci lay in 1D, along the contour of the chromosome). For instance, whereas HSPC loops shorter than 1Mb were bound by CTCF in 77.0% of cases (4.5-fold enrichment), loops longer than 3Mb were only bound by CTCF 46.7% of the time (a 2.9-fold enrichment) (Fig 1G). Similarly, longer loops were less likely to obey the convergent rule. For loops shorter than 1Mb, the CTCF motifs at loop anchors pointed inward 92.7% of the time, as compared to only 57.9% of the time for loops longer than 3Mb (Fig 1H).

When we visually inspected the Hi-C contact maps in HSPCs (*16*), we noted the presence of 408 additional long (>2Mb) loops that were not detected by our algorithms using the default parameters (Fig1D and E, FigS2a-d, Table S2) (*15*). Some of these loops were extremely large, spanning up to 117Mb (Fig 1F). The anchors of long loops exhibited minimal enrichment for CTCF (1.04-fold, Fig. 1G), and, even when CTCF was bound, they did not obey the convergent rule (130 of 290 CTCF motifs pointed inward) (Fig. 1H).

Taken together, these findings suggest that long loops form in HSPCs by a mechanism that is independent of CTCF.

Next, we sought to determine the basis of these long loops. We therefore examined the relationship between the long loops and DNA methylation canyons (Table S3). Specifically, we compared the position of loop anchors with the 282 canyons longer than 7.5kb (dubbed “grand canyons”), reasoning that shorter canyons might not reliably influence our HSPC contact map given the limitations on Hi-C resolution. Strikingly, we found that the rate of overlap with grand canyons depended strongly on the size of the loop. Of the anchors of HSPC loops shorter than 1Mb, 1.8% overlapped a grand canyon, representing a 4.4-fold enrichment (77 of 4287). By contrast, when we examined the anchors of HSPC loops longer than 3Mb that were annotated by HiCCUPS, we found that 29.0% overlapped a grand canyon – a 45-fold enrichment (18 of 62). Similarly, when we examined the 458 anchors of long loops identified by visual inspection, 24% (110) overlapped a grand canyon, a 16.7-fold enrichment (Fig 2A, B, C) (Fig 2D). (The reduced enrichment is likely due to the fact that the anchors of loops identified by visual inspection cannot be localized as precisely.)

**Figure 2.**
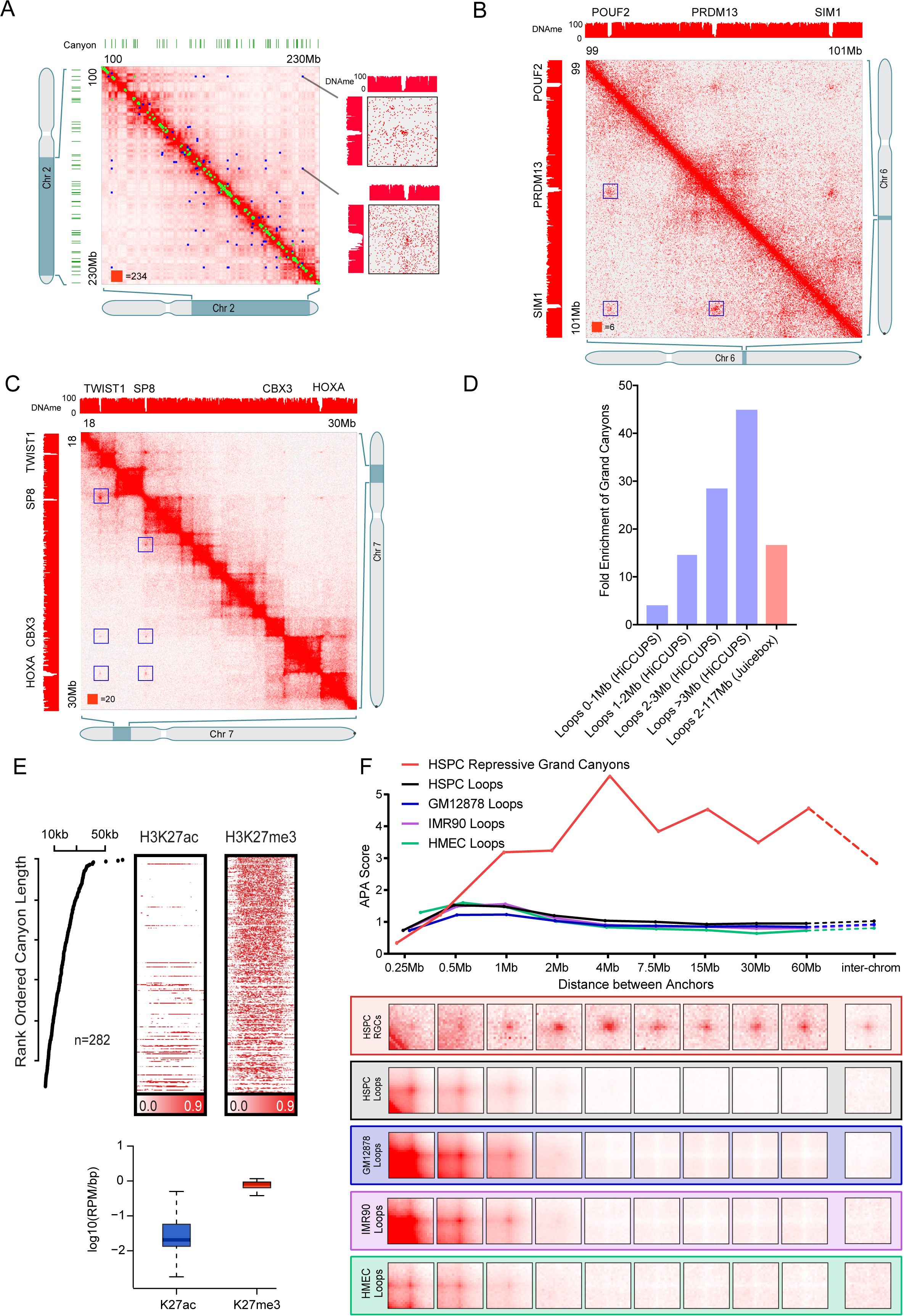
DNA methylation canyons and long-range interactions. **(A)** Example of HiCCUPS loops and Long loops on chromosome 2. Blue dots represent Long loops and green dots represent HiCCUPS loops. The matrices are shown at 100kb resolution and blow out of *POU3F3*, *HOXD* and *PAX3* regions at 5kb resolution. Green bars represent DNA methylation canyons. **(B)** Example of Long loops on chromosome 6 convergent on grand canyon regions. The matrices are shown at 100kb resolution and blowout of *POUF2*, *PRDM13* and *SIM1* regions at 5kb resolution. **(C)** Example of Long loops on chromosome 7 with grand canyons. The matrices are shown at 100kb resolution of *TWIST1*, *SP8* and *HOXA* regions at 25kb resolution. **(D)** Fold enrichment of Grand Canyons at loop anchors (HiCCUPS loops - blue bars, Long loops – red bar). **(E)** Strong enrichment of H3K27me3 and depletion of H3K27ac peaks in Grand canyons. **(F)** The aggregated peak analysis (APA) on Grand canyon interactions, with different length scale and inter-chromosomal interactions in HSPCs. Loop interactions are shown as a control. HMEC: Human Primary Epithelial Mammary Cells.

We were curious whether the histones at the anchors of long loops also exhibited particular epigenetic modifications. To probe this question, we performed ChIP-Seq in HSPCs using antibodies for histone marks indicative of repressive and active chromatin, specifically H3K27me3 and H3K27-acetylation (H2K27ac) respectively. We found that nearly all grand canyons (85%, 241 of 282) exhibit broad H3K27 trimethylation across the entire canyon interval, indicating a repressed state (Fig 2E). A small number lacked this mark, and instead exhibited broad H3K27 acetylation, indicating a more active state (18%, 53 of 282; note that, consistent with prior studies (*12*), nearly all grand canyons exhibited one of the two modifications). Crucially, the active, H3K27ac grand canyons were much less likely to be found at the anchors of long loops relative to grand canyons marked by H3K27me3 (5 of 39 vs 100 of 227, a 3.4-fold depletion) (Table S3).

Because the loops forming between grand canyons were so long, we wondered whether they can form links even when they lie on different chromosomes. To probe this question, we used Aggregate Peak Analysis (APA), a computational strategy in which the Hi-C submatrices from the vicinity of multiple putative loops are superimposed (*3, 16*). The enhanced contact frequency between pairs of loop anchors in aggregate is often visible via APA even when individual loops might not be discernable in a given map. This enhancement is indicated by an enrichment in the contact frequency at the center of the APA plot, and is reflected by an APA score >1.

Using APA, we examined pairs of grand canyons separated by varying distances along the contour of the chromosome, as well as pairs lying on different chromosomes. As controls, we examined the anchors of short loops we annotated in HSPC, as well as the anchors of loops annotated in prior *in situ* Hi-C experiments.

We found that pairs of grand canyons exhibit a tendency to form loops regardless of the linear distance separating them, and to form links even when located on different chromosomes (Fig 2F). By contrast, short HSPC loop anchors, and loop anchors annotated in earlier studies, showed enhanced proximity to one another only when lying on the same chromosome, at distances shorter than 2Mb (Fig 2F).

Next, we considered whether the long loops associated with repressive grand canyons that we had annotated in HSPCs were present in other human cell types (Fig 3A). We used APA to re-analyze a total of ~30 billion Hi-C read pairs from the 19 human cell types types in which loop-resolution Hi-C maps are available (*3, 8, 17–20*). In every cell type examined, the aggregate signal was either absent (18 cell types), or nearly absent (GM12878 B lymphoblastoid, where it was diminished by 88%) (Fig 3A,B; Fig S3). By contrast, loops associated with convergent CTCF sites in HSPCs were well preserved across cell types. The sole exception was a Hi-C map from HCT-116 cells in which cohesin had been degraded, a case where CTCF-mediated loops are now known to disappear (*20*). Visual examination of many long loops (*16*) was consistent with the above findings.

**Figure 3.**
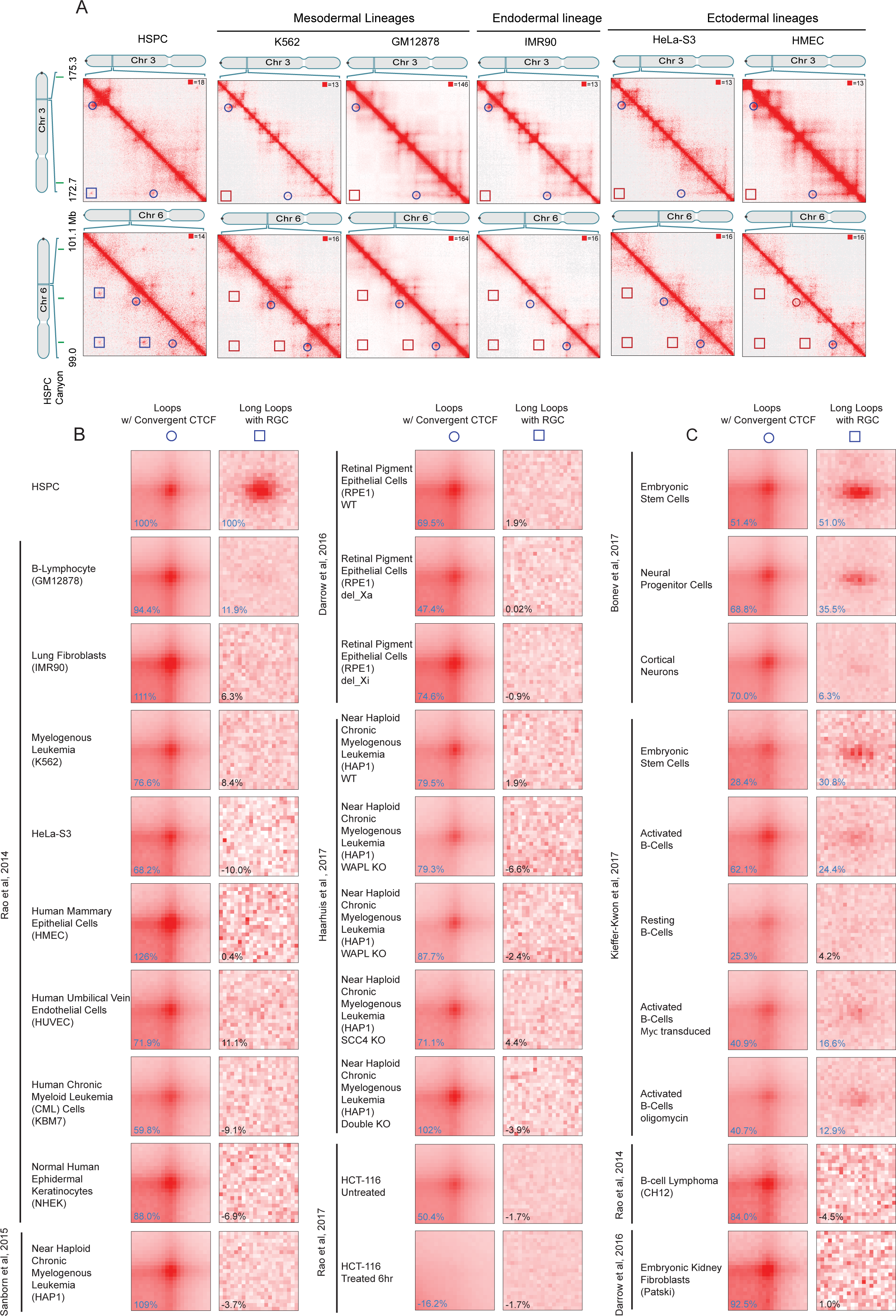
Canyon interactions are strongly enriched in undifferentiated cell types. **(A)** Example of HiCCUPS loops (circles) and Long loops (squares) on chromosomes 3 and 6 across cell types (left). APA for loops with convergent CTCF and Long loops with repressive grand canyons across cell types (right). Canyons are indicated in green (top). **(B)** APA on the indicated human cell types. **(C)** APA on the indicated mouse cell types.

We also used APA to analyze all 10 murine cell types in which loop-resolution Hi-C maps are available (>35 billion read pairs) (*17, 21*) to see if the long loops associated with repressive grand canyons that we had observed in HSPCs were conserved in mouse. We observed strong conservation in mouse embryonic stem cells and neural progenitor cells (*17*). All other more differentiated cell types had little or no discernable APA signal, with the exception of activated (but not resting) B-cells (Fig 3C, Fig S3). Together, these data suggest that long loops at repressed grand canyons are frequently seen in the stem and progenitor cell state.

We then examined this possibility further within more differentiated human hematopoietic lineages. We performed *in situ* Hi-C in two differentiated hematopoietic cell types, erythroid progenitor cells (EP cells, 857M contacts) and T-cells (622M contacts; see Fig 1A, Fig S4A-D). In both of these lineages, many of long loops with grand canyons were no longer visible (Fig 4A,B,C, Fig S5). To determine whether DNA methylation changes could account for this loss of long loops we examined DNA methylation data from both EP cells and T-cells. These analyses confirmed that canyons are preserved across all three cell types (Fig 4C), with modest changes in DNA methylation. Thus, these findings are consistent with a model where grand canyons tend to co-segregate in undifferentiated cells, such as HSPCs. This trend is also observed in the murine cells, with activated B cells as an exception (Fig S3).

**Figure 4.**
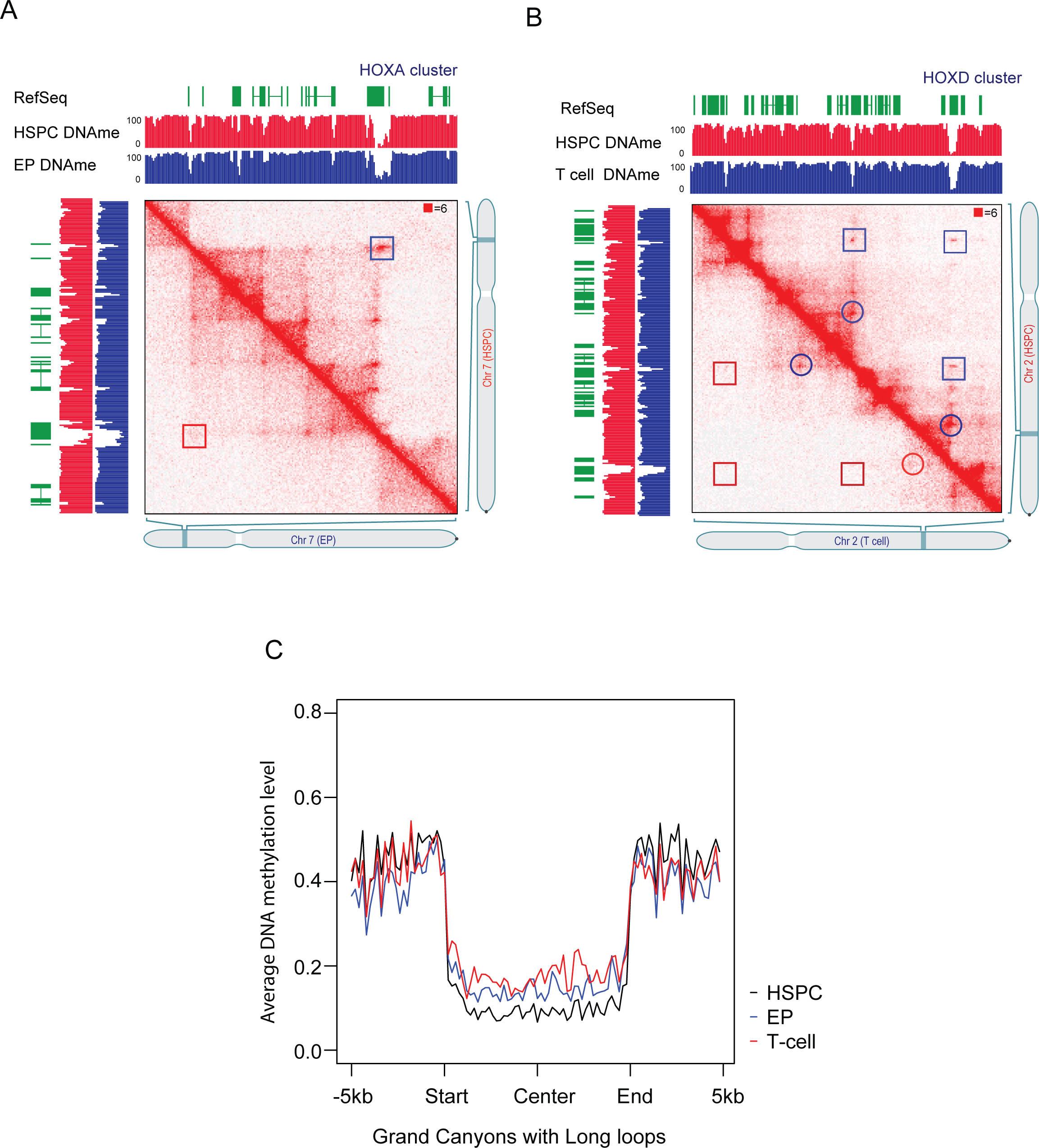
HSPC-specific Canyon interactions. **(A)** The comparison of HiCCUPS loops (circles) and Long loops (squares) between HSPC (upper) and Erythroid Progenitor (lower) on the *HOXA* cluster. DNA methylation in the HSPC and EP cells is shown. **(B)** The comparison of HiCCUPS loops (circles) and Long loops (squares) between HSPC (upper) and T-cell (lower) on *HOXD* cluster. DNA methylation in the HSPC and EP cells is shown. **(C)** The comparison of DNA methylation levels in HSPC, T-cells, and EP on Long loops overlapping grand canyon regions.

Finally, we sought to explore the functional significance of long loops by removing a grand canyon at a loop anchor. As almost all grand canyons are associated with promoters and exons, the functional impact of canyon deletion could be confounded by the effect of removing the gene. To obviate this concern, we identified a grand canyon that lay at the anchor of a long loop and contained no genes (“geneless” canyon, or “GLS”). GLS is 17 Kb long, lies 1.4 Mb upstream of the *HOXA1* gene, and forms long loops with a 28kb grand canyon in the *HOXA* region (Fig 5A,B,C, Fig S6A). GLS is a typical grand canyon, as it is coated with the repressive H3K27me3 mark, and has no transcriptional activity or active histone marks, confirming it does not serve as an enhancer (Fig 5B).

**Figure 5.**
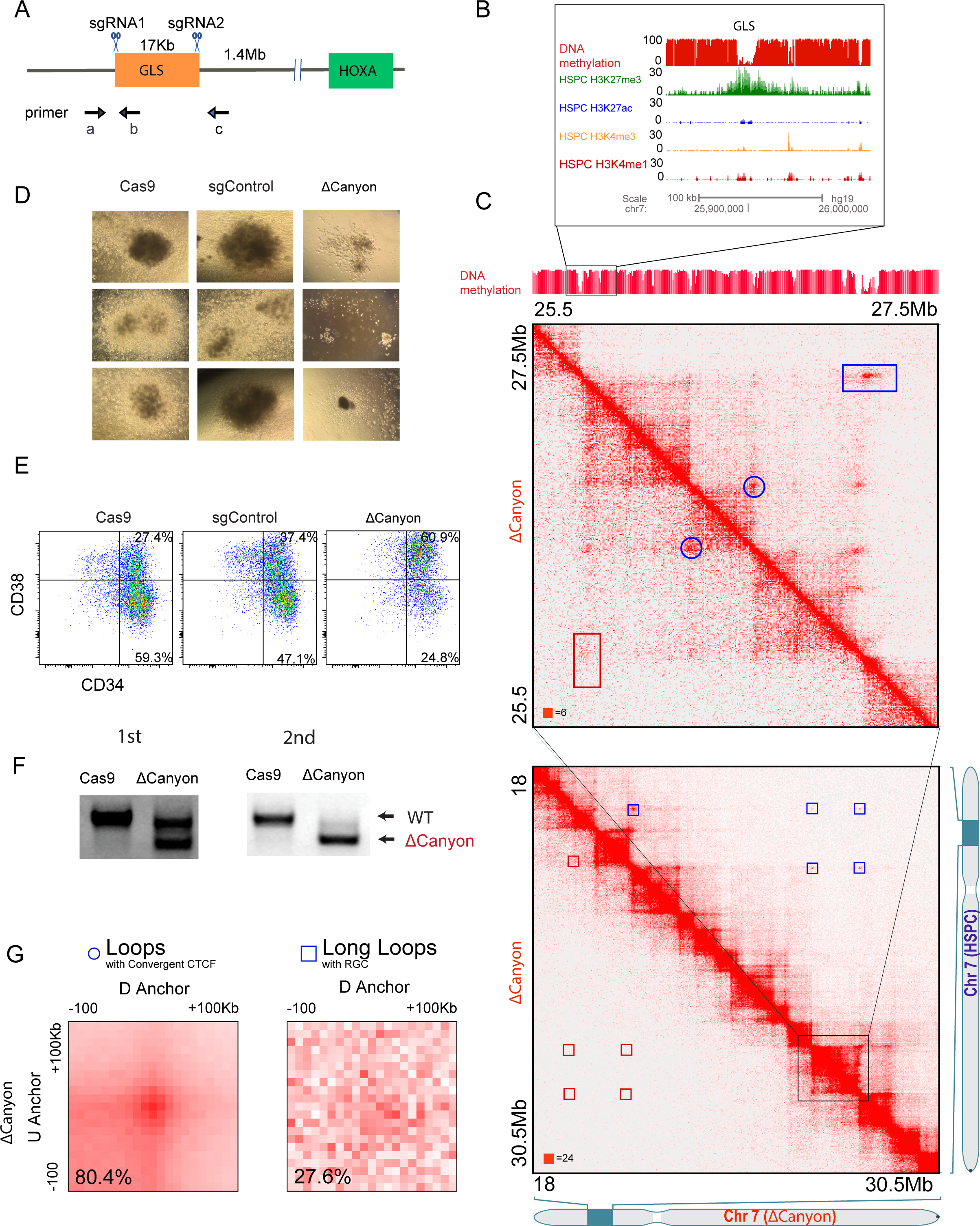
*HOXA* long-range Interactions Maintain HSPC Identity. **(A)** Schematic representation of CRISPR/Cas9 targeting of the geneless (*GLS*) canyon. **(B)** Epigenome browser track image of the geneless (*GLS*) canyon between the *MIR148A* and the *RNU6-16P* locus. H3K4me3, H3K27me3, H3K27ac, H3K4me1 and DNA methylation are displayed for HSPCs **(C)** Contact matrices of *GLS* and *HOXA* cluster region on chromosome 7 at 5kb resolution in WT CD34+ HSPCs (Upper) and canyon-deleted HSPCs (Lower) and Zoomed out contact matrices of the region including the *HOXA* cluster, *TWIST1* and *SP8* on chromosome 7 at 50kb resolution in WT CD34+ HSPCs (Upper) and canyon-deleted HSPCs (Lower). Circles: HiCCUPS loops, squares: Long loops. **(D)** Methylcellulose colonies from CD34+ HSPC after electroporation with Cas9 protein only (left), sgRNA + Cas9 protein (left), and sgControl (center). Representative images (same magnification) are shown. **(E)** Flow cytometry analysis of HSPCs treated with Cas9-only, control guide RNA, or guides to delete the *GLS* (Canyon). **(F)** PCR analysis of canyon deletion efficiency after two rounds of deletion prior to Hi-C. **(G)** APA for loops with convergent CTCF and Long loops with repressive grand canyons after Canyon deletion.

We deleted the GLS in HSPCs using a strategy we recently developed for efficient Cas9-mediated editing in primary cells (*22*) and assayed the pool of edited cells for colony formation ability.

Strikingly, after treatment with guide RNAs designed to delete the entire GLS, the number of colonies and their size was greatly reduced as compared to control experiments using either random guide RNAs or electroporation only (Fig 5D). After *ex vivo* culture, we performed FACS analysis for HSPC (CD34) and differentiation (CD38) markers. The cells treated to delete the GLS overwhelmingly acquired the marker CD38, indicating that they had differentiated. By contrast, the control cells predominantly expressed HSPC markers (CD34+ CD38-) (Fig 5E). Similarly, *HOXA* gene expression – an indicator of HSPC function– was greatly diminished after GLS deletion but not in control cells (Figure S6B).

Next, we performed *in situ* Hi-C on the cells after GLS deletion (Fig 5F). Strikingly, not only was the GLS-to-*HOXA* loop absent, but loops between repressed grand canyons were greatly attenuated genome-wide (Fig 5G, Fig S6C), consistent with the loss of HSPC state. Taken together, these data suggest either that the deletion of GLS compromised the primitive features of HSPCs, but was tolerated in the differentiated cells, or that the deletion of GLS directly promoted differentiation.

In this study, we have shown that long loops form in hematopoietic stem and progenitor cells. These loops differ from most loops that have been observed in prior Hi-C experiments in several important respects: (i) they bind CTCF much less frequently; (ii) they do not respect the CTCF convergent rule; and (iii) their anchors form links at arbitrary distances and when lying on different chromosomes. In all of these respects, the long loops and links we report in this paper closely resemble the loops and links that have recently been reported between superenhancers (i.e., long H3K27 acetylated genomic intervals) (*12, 13*). However, they also differ from those features in two important ways: (i) the loops and links in HSPCs connect repressed DNA methylation canyons, rather than superenhancers; and (ii) they form in primary cells, under physiological conditions, rather than in a cell line that has been engineered to bring about an abnormal genome conformation.

The loops we identify also resemble the superloops observed on the inactive X chromosome, insofar as they are very large and anchored at pairs of loci with similar modifications of H3K27 (*3, 12*). Finally, our findings are also entirely consistent with a very recent study, which identified long loops in mouse ES cells that were anchored at loci decorated by H3K27me3 and polycomb complex 1 (PRC1), and diminished upon differentiation (*17*).

Recently, we proposed that some loops form via cohesin-associated extrusion, whereas others form by compartmentalization (*3, 8, 20*). A crucial difference between these two mechanisms is that loop extrusion can only generate links among pairs of loci on the same chromosome, whereas compartmentalization can also lead to links among loci on different chromosomes. Thus, our findings here suggest that the long loops between repressed grand canyons do not form by extrusion. Instead, our data are consistent with a model in which repressed grand canyons in HSPCs tend to co-segregate by forming a subcompartment in the nucleus. This co-segregation could be a consequence of phase separation (*23–26*) or other mechanisms.

It is interesting to compare this subcompartment to those identified in prior studies. Multiple Hi-C studies have reported that loci bearing H3K27me3 tend to co-segregate (*17, 27, 28*), forming a subcompartment that is sometimes called “B1” (*3*). Similarly, microscopy studies have documented the presence of Polycomb bodies, in organisms ranging from *Drosophila* to mammalian cells (*29–31*). However, Polycomb decorates a significant fraction of the genome (*32–34*), whereas the subcompartment identified in the present study appears to be much smaller. In particular, H3K27me3 DNA methylation grand canyons correspond to only 241 genomic intervals, and together span only 3.5Mb: roughly 0.1% of the genome.

It is also interesting to speculate as to the function of this subcompartment. At present, its association with the HSPC state leads us to postulate that it plays a role in ensuring appropriate gene expression of the master regulators of cell identity and function, such as *HOXA* loci, through mechanisms we do not yet understand.

Our findings also clarify the nature of genome compartmentalization. In the earliest Hi-C maps, it was apparent that long intervals of chromatin (>100kb) exhibiting similar broad-source histone modifications tend to co-localize in the nucleus (*3, 4, 12*). Similar compartmentalization patterns were subsequently observed by many groups. Here, we expand on these findings in three ways. First, we observe hundreds of examples of co-segregation of intervals as short as 7.5kb in primary human cells. It is possible that patterns of compartmentalization exist at still finer scales which we cannot yet resolve. Indeed, we are unable to rule out the possibility that the epigenetic state of chromatin, at all scales at which it can be meaningfully interrogated, is associated with nuclear localization. Second, we find that subcompartments are not universally present across all cell types, but instead can be cell-type specific. Finally, we find a strong association between DNA methylation and compartmentalization, suggesting that CpG methylation plays a hitherto unappreciated role in the 3D architecture of the genome.

## Author Contributions

Conceptualizations: X.Z, M.J, X.H., M.A.G., and E.L.A.; Methodology: M.J., X.Z., A.P.A., and I.B.; Analysis: M.S.S., X.H., and J.Z. and J.R.; Investigation: M.J, X.H., X.Z., H.J., E.H.; Writing & Editing: X.Z., M.J., X.H., E.L.A., J.Z and M.A.G.; Supervision: W.L., E.L.A. and M.A.G.

## Acknowledgements

We thank the Dan L Duncan Cancer Center (Baylor College of Medicine) Core Grant (CA125123) for supporting this study. This work was supported by NIH grants: HG007538 and CA193466 (to WL), DK092883 and CA183252 (to MAG) as well as CPRIT grants. MAG is a Sam Waxman Cancer Research Foundation Investigator. MJ was supported by 5T32DK60445. E.L.A. was supported by an NIH New Innovator Award (1DP2OD008540-01), an NSF Physics Frontiers Center Award (PHY-1427654, Center for Theoretical Biological Physics), the Welch Foundation (Q-1866), an NVIDIA Research Center Award, an IBM University Challenge Award, a Google Research Award, a Cancer Prevention Research Institute of Texas Scholar Award (R1304), a McNair Medical Institute Scholar Award, an NIH 4D Nucleome Grant U01HL130010, an NIH Encyclopedia of DNA Elements (ENCODE) Mapping Center Award UM1HG009375, and the President’s Early Career Award in Science and Engineering.

All contact maps reported here can be explored interactively via Juicebox at http://www.aidenlab.org/juicebox/

## Supplementary Materials

Materials and Methods

Figs. S1-S5

Table S1- S3

References (35 to 37)

